# The Viral AlphaFold Database of monomers and homodimers reveals conserved protein folds in viruses of bacteria, archaea, and eukaryotes

**DOI:** 10.1101/2025.05.14.653371

**Authors:** Roni Odai, Michèle Leemann, Tamim Al-Murad, Minhal Abdullah, Lena Shyrokova, Tanel Tenson, Vasili Hauryliuk, Janani Durairaj, Joana Pereira, Gemma C. Atkinson

## Abstract

Viruses are among the most abundant and genetically diverse entities on Earth, yet the functions and evolutionary origins of most viral proteins remain poorly understood. Their rapid evolution often obscures evolutionary relationships, making it difficult to assign functions using sequence-based methods alone. Although conservation of protein fold can reveal deep homologies undetectable by sequence comparison, viral proteins remain vastly underrepresented in structural databases, limiting our ability to explore them at the structural level. Here, we address this gap by clustering all unique viral sequences from the NCBI RefSeq database and predicting the structures of ∼27,000 representative proteins using AlphaFold2, creating a large-scale viral structural resource, the Viral AlphaFold Database (VAD). We uncover ∼10,000 proteins belonging to clusters that share folds across viruses infecting bacteria, archaea, and eukaryotes, revealing shared protein folds across diverse host-infecting viruses. We also predict oligomeric states using AlphaFold2-based homodimer modelling, alongside structural comparisons to the Protein Data Bank, providing valuable new data on the potential for viral proteins to oligomerise. We further reveal that large regions of the viral protein universe remain functionally dark and report the discovery and experimental validation of a previously uncharacterised antiviral toxin-antitoxin (TA) system. VAD is a resource that provides a foundation for exploring viral structure–function relationships, including ancient folds that shape viral interactions across all life. Predicted structures used in this study are available at data-sharing.atkinson-lab.com/vad/.

## Introduction

Modern deep learning structural prediction methods, such as AlphaFold and RosettaFold have revolutionized the ability to use structural information in research [1-3]. As well as the use of inferred folds to make predictions about function that can be tested experimentally, structure versus structure comparison methods allow the discovery of homology that is not detectable at the sequence level using tools such as FoldSeek [4] and Distance-matrix alignment (Dali) [5]. This can be especially informative for proteins of viruses, which evolve fast due to short generation times, large population sizes, high mutation rates, and host-virus arms races [6, 7]. The taxonomic domain of viruses includes both eukaryote-infecting viruses, and bacteriophages that infect bacteria and archaea. Despite these two classes of viruses being very different due to their host specialisations, common folds of proteins have been identified. The single jellyroll (SJR) is one of the most common capsid folds, and can be found in various phages and eukaryote-infecting viruses [8], while the capsid protein of bacteriophage HK97 has homologues in herpes viruses, adenoviruses, along with viruses of archaea and protists [8, 9]. Most recently, RNA ligase T-like phosphodiesterases of both phages and eukaryote infecting viruses have been found to contain the same fold [10].

The ability to search for homologous proteins and infer function at the fold level depends upon the availability of large and diverse protein structure databases. The Protein Data Bank (PDB) has been an essential resource for structural comparisons for decades, storing and organizing experimentally determined structures of proteins and complexes [11]. A massive increase in the numbers and diversity of publicly available structures came with the release of the EBI AlphaFoldDB [12], which includes predicted structures for most proteins in the UniProt database, vastly expanding the number of available protein structures beyond what is present in the PDB. Structural comparison of these millions of protein folds has enabled the construction of networks of structural communities, annotated with their functional ‘brightness’ or ‘darkness’, depending on how well annotated the biochemical and biological functions of those proteins are [13].

A significant portion of the known protein fold universe remains functionally dark, and this is likely an underestimate of the true extent of structural and functional dark matter. One group that remains underexplored in structural biology is viruses. Despite being regarded as the most abundant biological entity on Earth and being an immense reservoir of genetic diversity [14, 15], viruses have so far been excluded from the EBI AlphaFold database. This is primarily due to the problem of polyproteins. Polyproteins are comprised of multiple viral peptides that are translated as a single concatenated peptide and subsequently proteolytically cleaved into individual mature proteins using viral or host proteases [16, 17]. While these polyprotein structures complicate structural prediction, the potential benefits of uncovering viral protein structural diversity outweigh these challenges. Recent viral structure databases, such as the BFVD [18] and ViralZone [10] projects, have addressed this gap to a large extent. However, these resources rely on less accurate methods than the reference AlphaFold2 implementation and are limited to monomeric structural predictions.

Here we have clustered 647,000 unique viral sequences from the NCBI RefSeq database [19], predicted the structure of ∼27,000 representatives using AlphaFold2 [1], and predicted higher order oligomeric states where possible. This represents a high-quality resource for exploring viral protein diversity, including in the context of host taxonomy. We have identified 1,142 clusters of viral proteins that share the same fold across viruses infecting two or more domains of cellular life (bacteria, eukaryotes, or archaea). We have characterized functional darkness of viral folds, predicted homodimerization tendencies and structures for almost all proteins in our database, and predicted higher-order multimeric states where possible. Through searching for functionally dark protein structures in conserved neighbourhoods, we have uncovered and experimentally validated a new Type II toxin–antitoxin (TA) system, KreTA, found in prophages of bacteria, and mapped conserved and variable folds across virus–host interactions, including defence and anti-defence systems.

## Results and Discussion

### Clustering the diversity of viral proteins at the sequence level reveals homology among proteins from viruses that infect hosts in different domains of life

Due to the computational challenges of predicting structures for all available viral protein sequences, we aimed to assemble and fold a representative subset that captures broad diversity across sequence space. To do this, we first downloaded protein sequences from the NCBI RefSeq database [19], limiting by taxonomy to virus (NCBI taxid 10239). The taxonomic distribution of these 647,000 sequences is depicted in **Fig. S1**, spanning the ten most abundant viral taxa across the kingdom, phylum, and family levels. The host of each virus was assigned by querying the Virus-Host database [20]. Protein sequences were then clustered using MMseqs2 [21], with relatively lenient identity and coverage thresholds (30% and 0.01 respectively) to minimize the number of clusters while maintaining sufficient separation between them (**Fig. 1A**). This clustering process yielded 117,479 clusters and 61,868 singletons, with an average cluster size of 5.5 members (**Table S1, Dataset S1**).

**Fig. 1.**
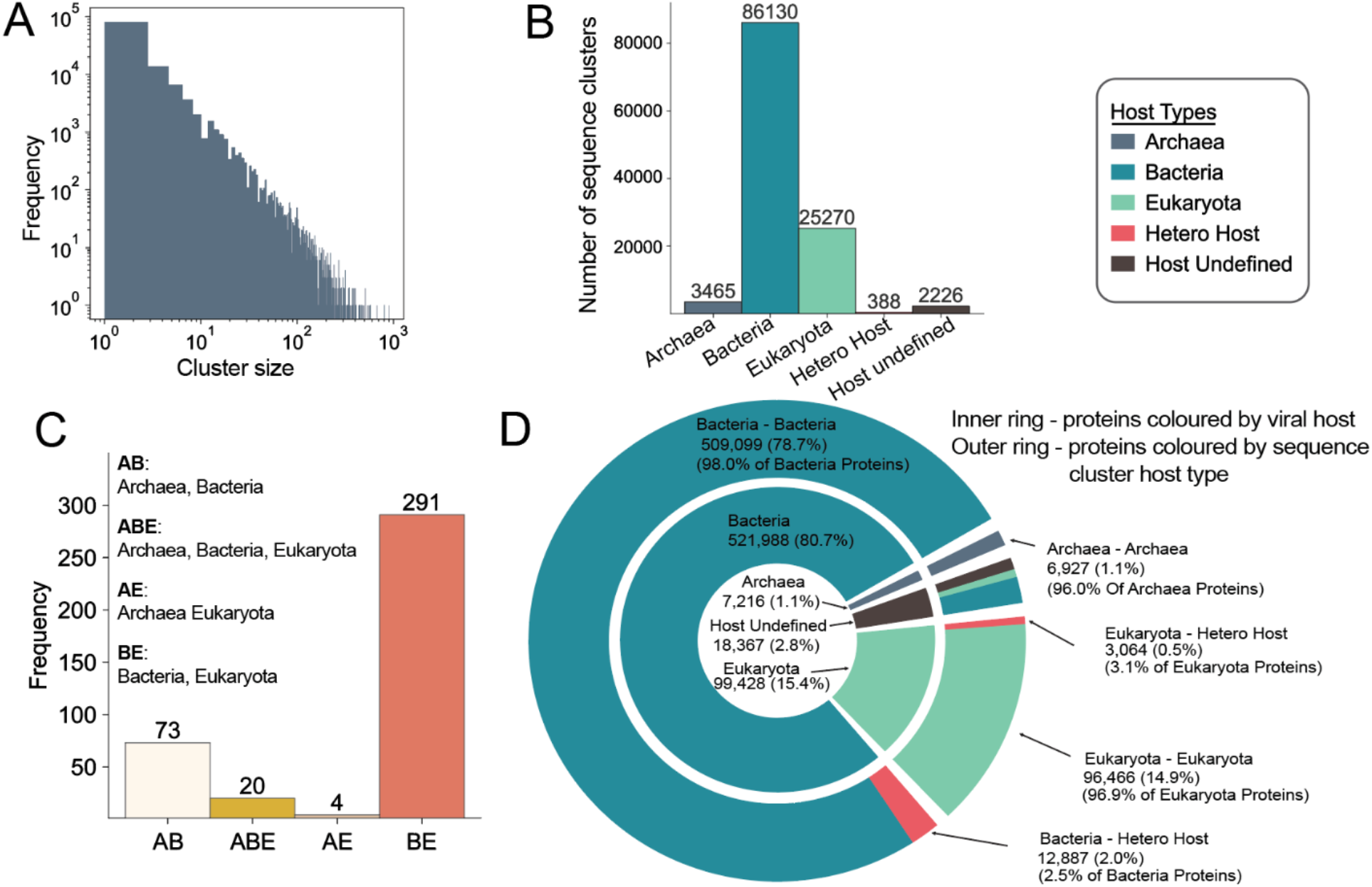
Sequence clustering and host-type assignment of viral proteins and clusters. (**A**) Distribution of sequence clusters by size. (**B**) Distribution of sequence clusters by host type. (**C**) Distribution of sequence hetero-host clusters based on host domain composition. (**D**) Distribution of proteins by viral host (inner ring) and sequence cluster host type (outer ring), where viral host segments correspond to cluster type segments, displaying the cluster host type membership of proteins by viral host.

The assignments from the Virus-Host database [20] allowed us to group proteins by host domain (bacteria, archaea or eukaryotes), where a cluster with multiple hosts associated with the proteins within is referred to as a hetero-host cluster. It is important to note that this does not imply that the same protein can be found in viruses infecting vastly different hosts; rather, the cluster is comprised of homologous proteins that can be found in different host-infecting viruses. Therefore, hetero-host cluster composition reflects molecular patterns that are common across viral life. Reflecting the original taxonomic distributions, most clusters comprise proteins from viruses infecting bacteria (**Fig. 1B**) In total, 388 clusters were assigned a sequence-level hetero-host label, most of which comprise proteins from viruses with either bacterial or eukaryotic hosts (**Fig. 1C**). The distribution of host domain (eukaryota, bacteria or archaea) before and after clustering is shown in **Fig. 1D**. To reduce ambiguity, we excluded from these counts any viruses annotated as infecting other viruses, such as virophages that parasitize giant viruses, and phage satellites [22] . Most unique viral protein hosts are bacterial (79.9%), followed by eukaryotes (15.4%). Archaeal hosts are assigned for only 1.1% of the proteins (**Fig. 1D**).

While some of the sequence-level clusters contain proteins annotated as structural proteins, i.e. tail fibers, capsids, or their chaperones, the majority of hetero-host clusters represent proteins that are annotated to interact with or modify nucleotides or nucleic acids (**Dataset S1**). The most prevalent are proteins involved in replication, nucleases, nucleotide hydrolases, nucleotide modification enzymes, helicases, nucleic acid ligases and methyltransferases. The largest hetero-host sequence cluster (567 proteins, found in viruses infecting either bacteria or eukaryotes) contains homologues of cytidine and deoxycytidylate deaminase –an enzyme which viruses can use to modulate nucleotide pools to favour their own replication [23], and sometimes serves as a component of antiphage defence [24]. The second biggest sequence cluster contains proteins homologous to the clamp loader of DNA polymerase, which enables efficient and processive replication and is found in bacteriophages and viruses of protists. Finally, there are 20 sequence clusters spanning bacteria, eukaryote, and archaeal hosts. The largest of these contains DNA methyltransferases (**Dataset S1**). Thus, at the sequence level, conserved nucleotide-processing functions are often the most readily identifiable across host domains.

### Prediction of representative protein structures to create the Viral AlphaFold Database (VAD)

Representatives from the viral protein sequence clusters were folded with AlphaFold2 [1], limiting to clusters with size five or greater. To sample sequence diversity in smaller clusters, we also randomly selected a set of 7,243 sequence clusters, with an average cluster size of 1.6. We did not set a strict amino acid length limit but rather folded all proteins that we could, given our computational resources. Compared to the two other predicted structure databases for viral proteins, BFVD [18] and ViralZone [10], our length distribution is most similar to BFVD, with an average length of 176 amino acids (**Fig. S2**). The length distribution in our database has a longer tail due to not using a length cut-off; our largest predicted fold is from a 3,595 amino acids long protein. ViralZone predictions tend to be longer on average, probably due to eukaryote-infecting viruses having longer proteins on average than phages. The final Viral AlphaFold Database (VAD) structure dataset consists of 26,962 predictions for monomers and 26,754 homodimer co-folding attempts (of which only a subset are confident homodimers; see below) (**Dataset S1**). The taxonomic distribution of VAD is depicted in **Fig. S3** (full lineage information is found **Dataset S1**), and covers the full diversity of viral kingdoms, phyla and families seen in the starting sequence set (**Fig. S1, S3**).

The quality of the predictions in the dataset is assessed with the pLDDT metric [1]. The VAD dataset has a global average pLDDT of 78, with the majority of predictions reaching an average pLDDT of approximately 90, reflecting high overall model confidence (**Fig. 2A**). Compared to ViralZone [10] and BFVD [18], VAD demonstrates improvements in prediction quality. Among near-identical proteins (>90% sequence identity and coverage) shared between VAD and BFVD, 74% of VAD models have higher average pLDDT scores, and 54% show an increase of more than 5 units. These results underscore the accuracy gains enabled by AlphaFold2’s more computationally intensive modelling, particularly in capturing reliable structures for diverse viral proteins.

**Fig. 2.**
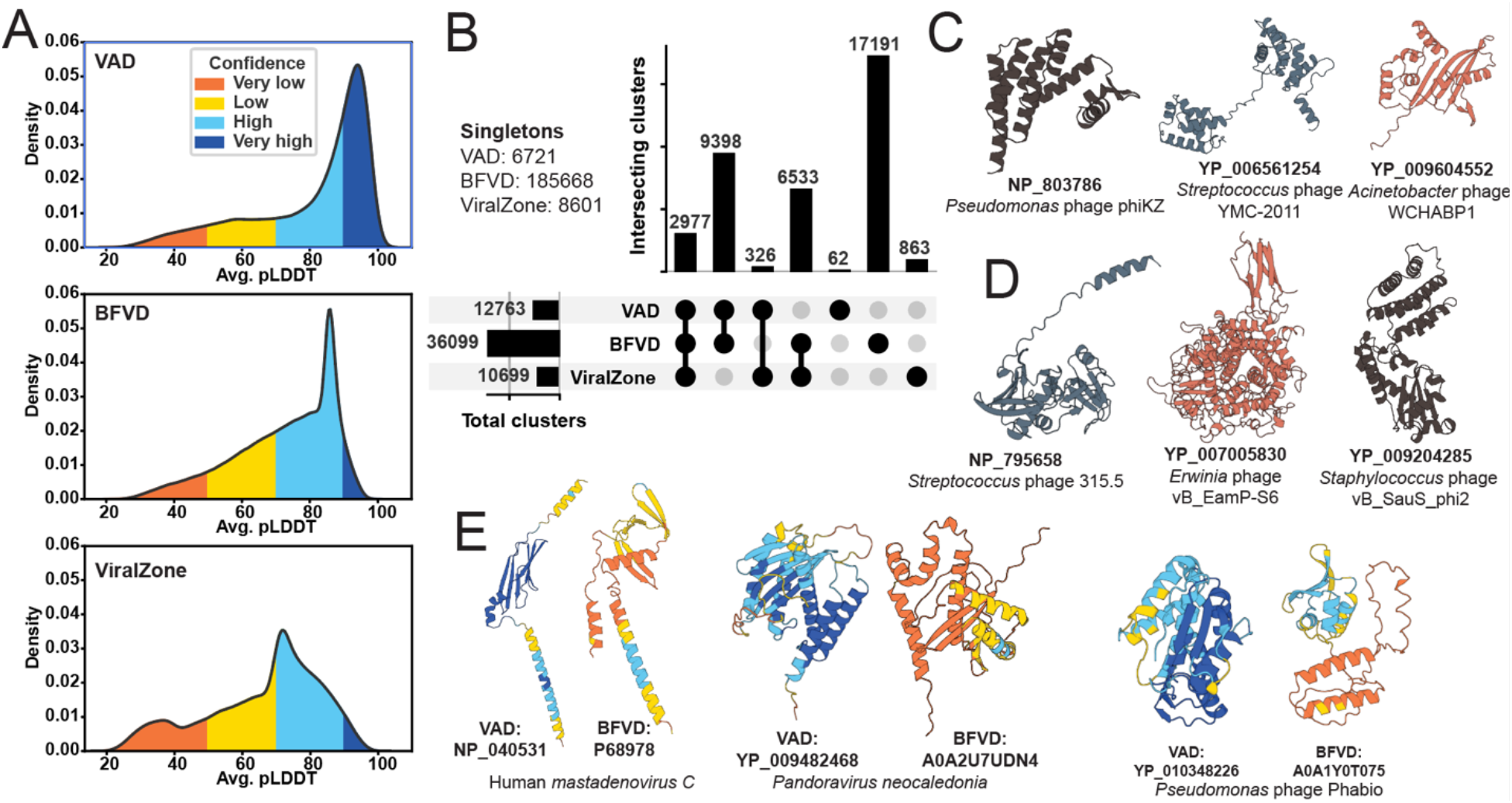
VAD comparison with other databases of viral predicted structures. (**A**) Density plot of Average pLDDT distributions for VAD, BFVD, and ViralZone. pLDDT is averaged per prediction in each dataset. (**B**) UpSet plot detailing overlaps of non-singleton structural clusters (with Foldseek easy-cluster coverage 0.7 and default parameters) across VAD, BFVD, and ViralZone with singleton cluster counts listed (**C**). Three examples of VAD cluster singletons with >80 pLDDT. (**D**). Three examples of clusters unique to VAD with >80 pLDDT. (**E**). Three examples where the VAD structure pLDDT is higher than the corresponding BFVD structure pLDDT.

Structure-based all-vs-all searches using the ∼27,000 VAD cluster proteins identify more than twice as many unique cluster-to-cluster relationships as sequence-based searches can find across all 647,000 viral proteins. This indicates that this relatively small, structure-representative set effectively captures relationships within the viral NCBI RefSeq database. VAD is structurally diverse, as seen by the 12,763 structural clusters and 6,721 singletons encompassing the 27k VAD cluster proteins (**Fig. 2B**). In addition, despite its smaller size, the VAD dataset spans a substantial portion of the non-singleton structural clusters found in both BFVD and ViralZone, which contain 2.5× and 13× more structures, respectively (**Fig. 2B**). Examples of singletons and unique VAD clusters are shown in **Fig. 2C and D** respectively. When comparing near-identical VAD and BFVD proteins (>90% identity and coverage), 74% of VAD proteins have a higher average pLDDT than the BFVD counterparts, with 54% having a difference of over 5 pLDDT units, indicating that the more computationally expensive AlphaFold2 modelling can have a big accuracy impact in predicting viral protein structures. Examples of such cases are shown in **Fig. 2E**. Together, these results show that the VAD dataset is both structurally diverse and high confidence, making it a valuable resource for structural searches and for advancing our understanding of viral protein function and evolution.

### Proteins with the same structural fold in VAD are found in viruses infecting bacteria, archaea, and eukaryotes

Sequence clustering shows that the vast majority of cluster representatives come from viruses infecting bacterial hosts (**Fig. 1B**). To test whether structure-based clustering reveals broader relationships across host types, we examined whether proteins from viruses infecting different domains of life share similar folds even when their sequences diverge. We therefore clustered the 26,962 structures in VAD using Foldseek [4] in TMalign mode, which groups proteins by structural similarity. This resulted in 12,894 clusters with 9,753 singletons, with an average cluster size of 2.1 (**Table S2, Dataset S1**). Host domains (bacteria, archaea, or eukaryotes) were then assigned to structural clusters using the same host annotations and criteria as for sequence clusters.

Structural clusters show a shift in host distribution: although bacterial host type clusters still dominate, the proportion of eukaryotic host type clusters increases (**Fig. 3A**). Hetero-host clusters also expand markedly; 913 clusters include proteins from both bacterial and eukaryotic viruses (BE), with additional clusters spanning archaea-bacteria (AB), archaea-eukaryota (AE), and all three domains (ABE) (**Fig. 3B**). While fewer than 1% of VAD proteins belong to hetero-host sequence clusters, this jumps to 35.8% in structural clusters (**Fig. 3C**). This striking increase suggests that folds are indeed conserved across greater evolutionary distances and are more frequently shared across viral life than sequence similarity alone would indicate.

**Fig. 3.**
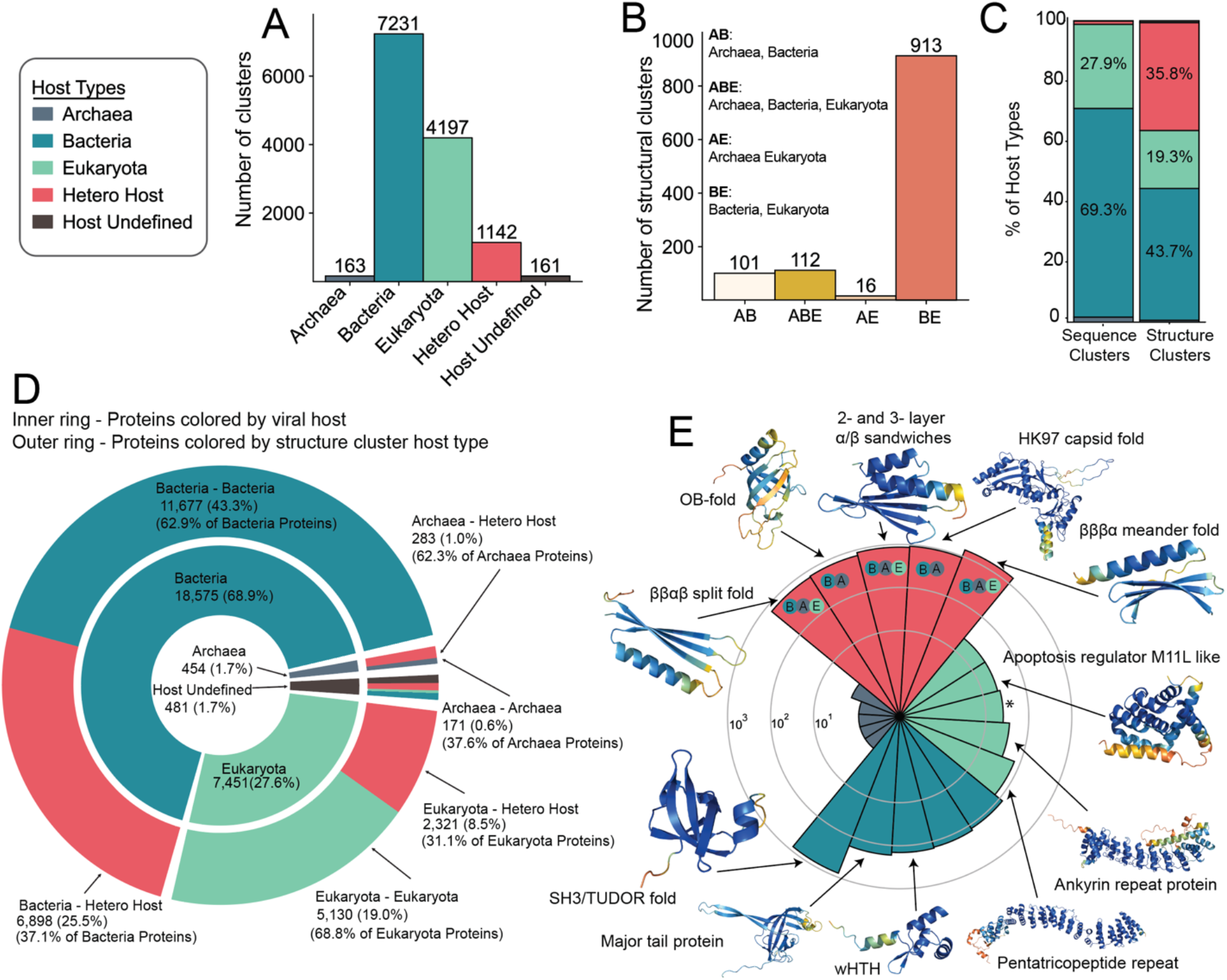
Structural clustering of VAD proteins reveals core folds in viruses infecting different domains of life. (**A**) Distribution of structural clusters by host type. (**B**) Distribution of structural hetero-host clusters based on host domain composition. (**C**) Sequence and structure cluster host-type distribution of VAD proteins. (**D**) Distribution of proteins by viral host (inner ring) and structural cluster host type (outer ring), where viral host segments correspond to cluster type segments, displaying the cluster host type membership of proteins by viral host. (**E**) Radial Histogram of the five largest structural clusters, their sizes and the structures of select cluster representatives colored by pLDDT. The contributing host taxa are shown within the hetero-host segments (B: bacteria; A: archaea; E: eukaryota). The third largest eukaryote specific cluster (15 proteins; marked with an asterisk) consists of a single helix topology and is not shown.

Next, we examined how proteins from viruses infecting specific host domains are distributed across structural clusters (**Fig. 3D**), as we did for sequence clusters in **Fig. 1D**. The majority of proteins from bacteriophages (62.9%) fall into clusters that also contain proteins from other bacterial viruses, but a substantial fraction (25.5%) are in hetero-host clusters. Similarly, 31.1% of proteins from viruses infecting eukaryotes are in hetero-host clusters, suggesting widespread structural reuse across viral lineages despite divergent host range. Archaeal viruses also show notable structural overlaps, with 62.3% of their proteins falling into archaeal-only clusters and 37.6% into hetero-host clusters.

To identify common structural themes across diverse viruses, we examined the largest structural clusters in more detail (**Fig. 3D**). Most large structural hetero-host cluster representatives are small mixed α/β folds, comprising ∼4% of all proteins belonging to structural hetero-host clusters (**Dataset S1**). The largest, third, and fifth largest cluster representatives are α-β sandwiches: two meander folds and one split fold. The fourth largest cluster representative is an α-β OB-fold (**Fig. 3E**). These small folds are typical ‘*urfolds’* – 3D architectures that can be formed from vastly different sequences, allowing for extensive functional innovation [25]. The second largest structural hetero-host cluster representative features an HK97 fold found in capsid proteins of various viruses. Overall, apart from the capsid fold, the largest clusters seem to represent common ancient core folds, not limited to viruses but also found in proteins across all life on Earth.

Most of the large structural eukaryotic mono-host cluster representatives are α-domain structures. The two largest of these clusters are likely pentatricopeptide and ankyrin-repeat proteins, where both repeat four-helix bundles. These tandem repeat clusters comprise ∼1% of all proteins belonging to structural eukaryotic clusters and ∼3% of proteins belonging to non-singleton clusters (**Dataset S1**). The third largest cluster is a singular α-helix with unstructured termini; the members of this cluster are similarly simple, with a minority exhibiting a turn between two α-helices. The fourth largest of these clusters is a globular α-domain protein, the M11L-like apoptosis regulator (**Fig. 3F**). Thus, most of these eukaryote-host-specific protein folds are likely all involved in hallmark eukaryote-like protein-protein and protein-RNA interactions [26, 27]. This may reflect viral adaptation to the complex and compartmentalised regulation characteristic of eukaryotic hosts [28].

Bacterial mono-host cluster representatives consist mostly of small β-barrel folds. The largest, fourth, and fifth largest of these clusters have SH3/Tudor-like folds (**Fig. 3F**). These putative Tudor fold clusters cover ∼1.3% of all proteins in bacterial structural clusters and ∼2.6% of proteins belonging to non-singleton clusters (**Dataset S1**). The second largest cluster contains tail tube proteins from λ-like phage tail tube proteins, while the third largest cluster contains winged helix-turn-helix (wHTH) folds that are often involved in DNA binding [29] (**Fig. 3F**). Archaeal structural clusters consist almost entirely of singletons, where member proteins are uncharacterized, reflecting the paucity of data for viruses infecting this domain of life.

### Oligomerisation states across viral proteomes

Many viral proteins exert their biological functions as oligomers rather than monomers. To explore oligomerisation tendencies, we predicted homodimeric structures for all but the ∼200 largest proteins of VAD monomers. These homodimers include 26,754 predictions, of which 11,313 exhibit a confident fold and 2,957 exhibit a good fold (with ipTM thresholds set to 0.5 for “confident” and to 0.8 for “good”) (**Fig. 4A**). Multiple VAD monomers do not form homodimers; in fact, the average ipTM score for VAD homodimers is 0.44 (**Dataset S1**), below the cut-off for “good” models. In order to determine if low-quality monomeric predictions skewed the quality of homodimer counterparts, we filtered monomers with plDDT scores lower than 50 (**Fig. S4**). The effect of this filtering was marginal without much change in the distribution density of predictions by ipTM score. In 1188 cases, the pLDDT of the dimer (max across the 2 chains) is >5 units higher than the monomer pLDDT. **Fig. 4B** shows three extreme examples, where the monomer and dimer structures have significant differences. Where we do see good scores for homodimerization, this can include homodimers in the strictest sense (two monomers interacting alone), as well as larger homo and heterooligomeric complexes that include an interface between two identical protein subunits. To address potentially larger complexes of VAD proteins we predicted oligomeric arrangements of monomers in VAD. We compared monomeric structures with those in the PDB: structural hits of monomers found more than once in a complex in a PDB structure were classified as homo-oligomeric (3,864 VAD clusters), while those found with other monomers were classified as hetero-oligomeric (2,623 VAD clusters) (**Dataset S1**). **Fig. 4A**, turquoise density, shows the dimer iPTM distribution of all VAD clusters compared to those having homomer PDB hits, showcasing a shift towards higher iPTMs for the latter. Importantly, AlphaFold-multimer seems able to predict dimeric structures in their higher order oligomeric state, as shown by the two examples in **Fig. 4C** where the closest Foldseek-Multimer PDB hit is a homo-trimer and the predicted dimer adopts the trimeric configuration. As AlphaFold-Multimer was trained on complexes from the PDB, predicted oligomeric conformations may simply reproduce those present in the training data for close homologues. In addition, PDB structures used as templates can project oligomeric states onto homodimeric predictions. However, oligomeric awareness can also be seen at low sequence identities, as seen in the dUTPase examples in **Fig. 4C**, which share only 28% sequence identity.

**Fig. 4.**
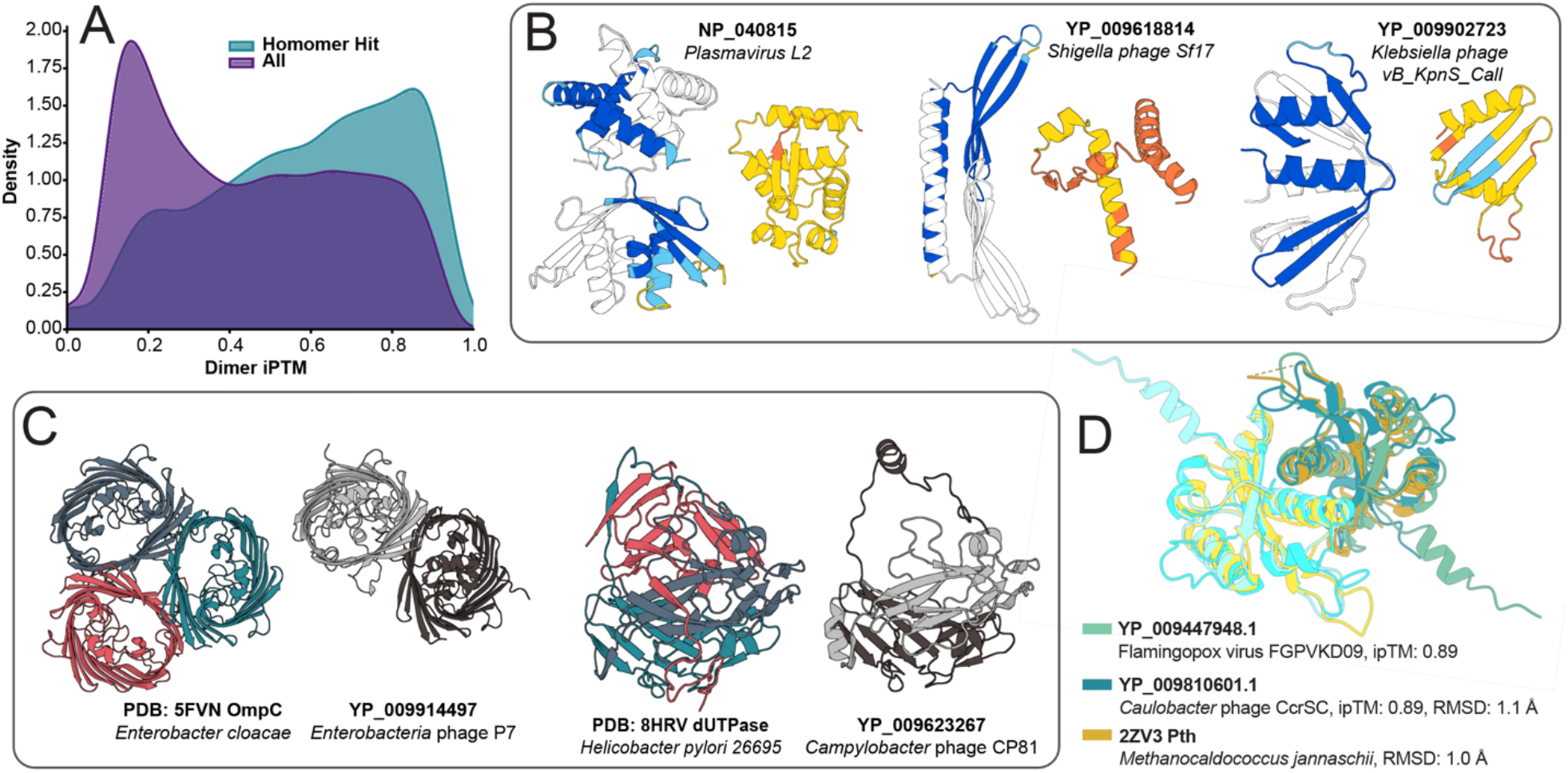
A large proportion of proteins in VAD are predicted to form homodimeric interfaces. (**A**) Density plot of ipTM distribution for all VAD proteins, and those with a homomer (homo-oligomer or homodimer) hit in the PDB. (**B**) Cases where the prediction quality drastically increases in the homodimer compared to the monomer. (**C**) Examples of cases where VAD homodimers are found in oligomer-aware conformations. Pairwise sequence identities are 65% in the case of the OmpC homologues and 28% in the case of the dUTPases. (**D**) Confident dimer predictions of Pth homologues from viruses that infect both bacteria and eukaryotes

One surprising observation in VAD is the prevalence of peptidyl tRNA hydrolases (Pth) across a variety of different viruses and phages. By cleaving peptides bound to tRNA, Pth plays a crucial role in protein synthesis by both rescuing stalled ribosomes and freeing up the pool of free tRNAs [30, 31]. We have found that Pth is encoded in various bacteriophages and viruses that infect birds and insects, and its dimeric structure is conserved (**Fig. 4D**). A functional Pth encoded in the bacterial host has previously been found to be important for λ phage translation of two-codon minigenes that are otherwise toxic due to accumulation of ‘dropped-off’ peptidyl-tRNA [32]. Thus, by carrying their own Pths, viruses may ensure efficient translation of their proteins and avoid ribosome stalling and toxicity. Our predictions suggest that the Pth sequence level hetero-host clusters are primarily homodimers. Superposition of AlphaFold Pth predictions from Caulobacter phage and Flamingopox virus with the PDB structure of Pth from archaeon *Methanocaldococcus jannaschii* show excellent structural alignment (**Fig. 4D**).

### Structural matches to the Protein Atlas; functional darkness and brightness of viral protein structures

We asked how similar or dissimilar viral proteins are to other proteins with known structures. A Foldseek search of VAD structures against UniProt3D community representatives and the AFDB50 database revealed 12,473 representatives (covering 292,376 viral sequences) falling into connected components in UniProt3D, 209 representatives (3,635 sequences) into unconnected “dust” UniRef50 clusters, 2,627 representatives (55,922 sequences) having matches to AFDB proteins not present in UniProt3D v1 due to the pLDDT threshold of 90 used, and 9,279 representatives (115,750 sequences) having no good match to either database (**Fig. 5A**). Among those with matches to known structural communities, most are functionally bright— a term used to describe proteins with known or inferred biological function [13]. However, a substantial fraction falls into dark communities, meaning they lack functional annotation and remain poorly understood despite having a predicted structure (**Fig. 5B**). When examined by host taxonomy, proteins from viruses infecting bacteria and archaea were more likely to have high-confidence structural matches to the AFDB, whereas proteins from eukaryotic viruses had the highest proportion of no matches (**Fig. 5C,D**). Among proteins that did yield a structural hit, the confidence (as measured by LDDT) was broadly similar across host types, suggesting that once a structural analogue is found, the quality of the alignment does not depend strongly on host taxonomy.

**Fig. 5.**
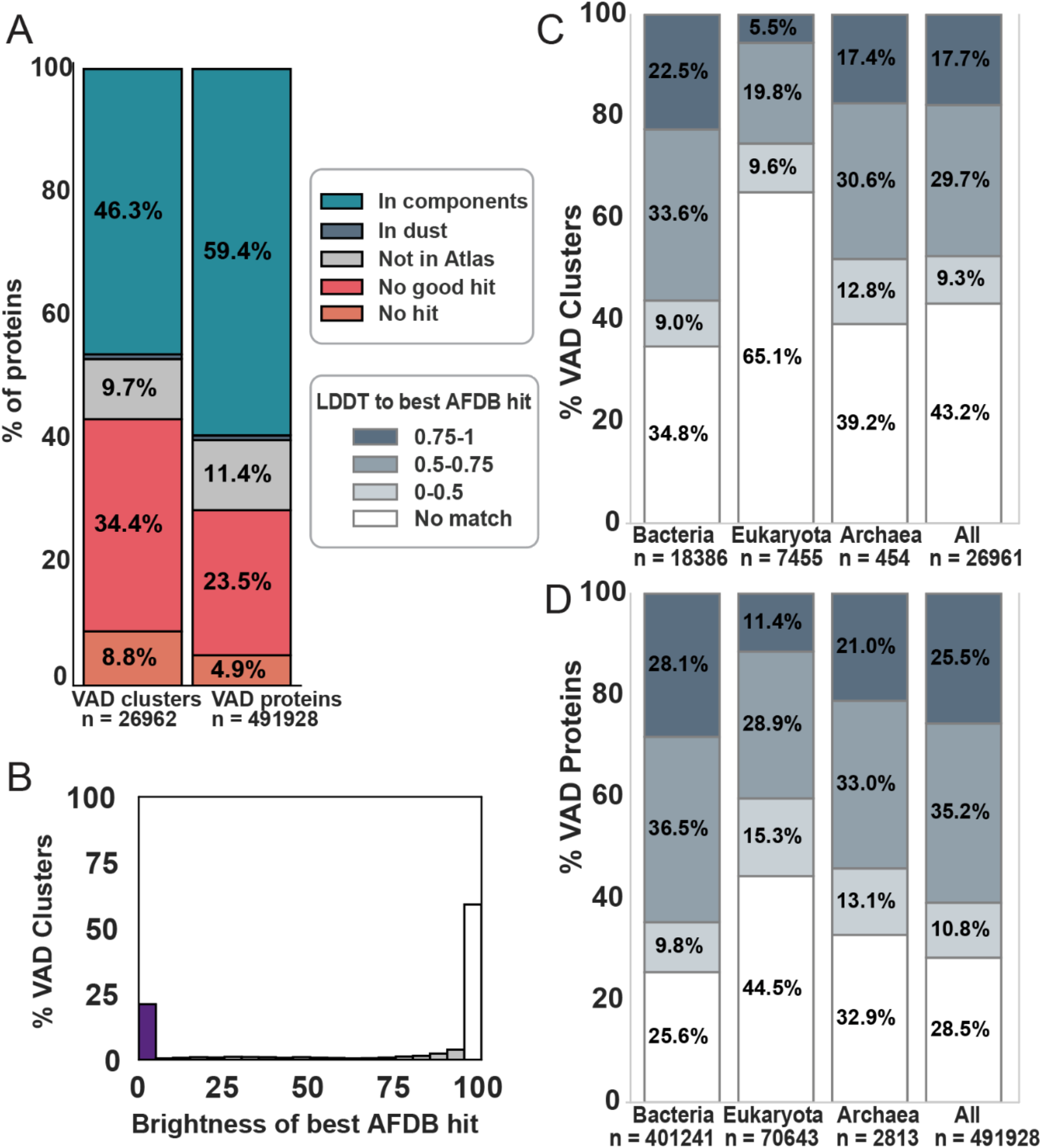
Structural similarity of VAD to the AlphaFold database. (**A**) Division of VAD representatives and proteins into those without any hits to AFDB50 or UniProt3D, without hits with TM-score > 0.5 without hits to UniProt3D (i.e only hits AFDB structures with pLDDT < 90), hits only to dust, and hits to connected components in UniProt3D. ‘Representatives’ mean cluster representatives that we have predicted the structure for, and therefore are present in VAD. ‘Proteins’ extends the count to all proteins in the clusters. (**B**) Distribution of functional brightness values for the brightest AFDB hit for each VAD representative. (**C**) LDDT of best hits across different host superkingdoms as per lower inset box (**D**) LDDT of best hits across different host superkingdoms expanded to proteins in each representative cluster.

### VAD proteins are related to functionally dark bacterial protein communities that include prophage-encoded toxin-antitoxin systems

A previous exploration of the universe of functionally dark protein folds led to the discovery of new Type II toxin-antitoxin (TA) systems [13]. Since we have uncovered additional dark proteins in VAD, we asked whether there could be further undiscovered TAs in our data. We focused on VAD proteins in Protein Atlas communities that also include bacteria, since TAs can be frequently found on prophages integrated into bacterial chromosomes [33, 34]. Specifically, we focused on small-operon-encoded proteins encoded in communities containing *E. coli*, the model system for our validations. Using GCsnap and FlaGs we analysed gene neighbourhoods, searching for conserved bi-cistronic gene operons characteristic of toxin-antitoxin systems. We found a pair with homologues across Gammaproteobacteria, consistently localized in prophage-like regions of *Enterobacteriales* (**Fig. 6A**). This includes annotated pathogenic strains isolated from clinical and food sources. The conserved two gene architecture is strongly suggestive of a TA system. We name this putative TA system KreTA after the mythical heroic twins in the Albanian folk epic *Kângë Kreshnikësh*.

**Fig. 6.**
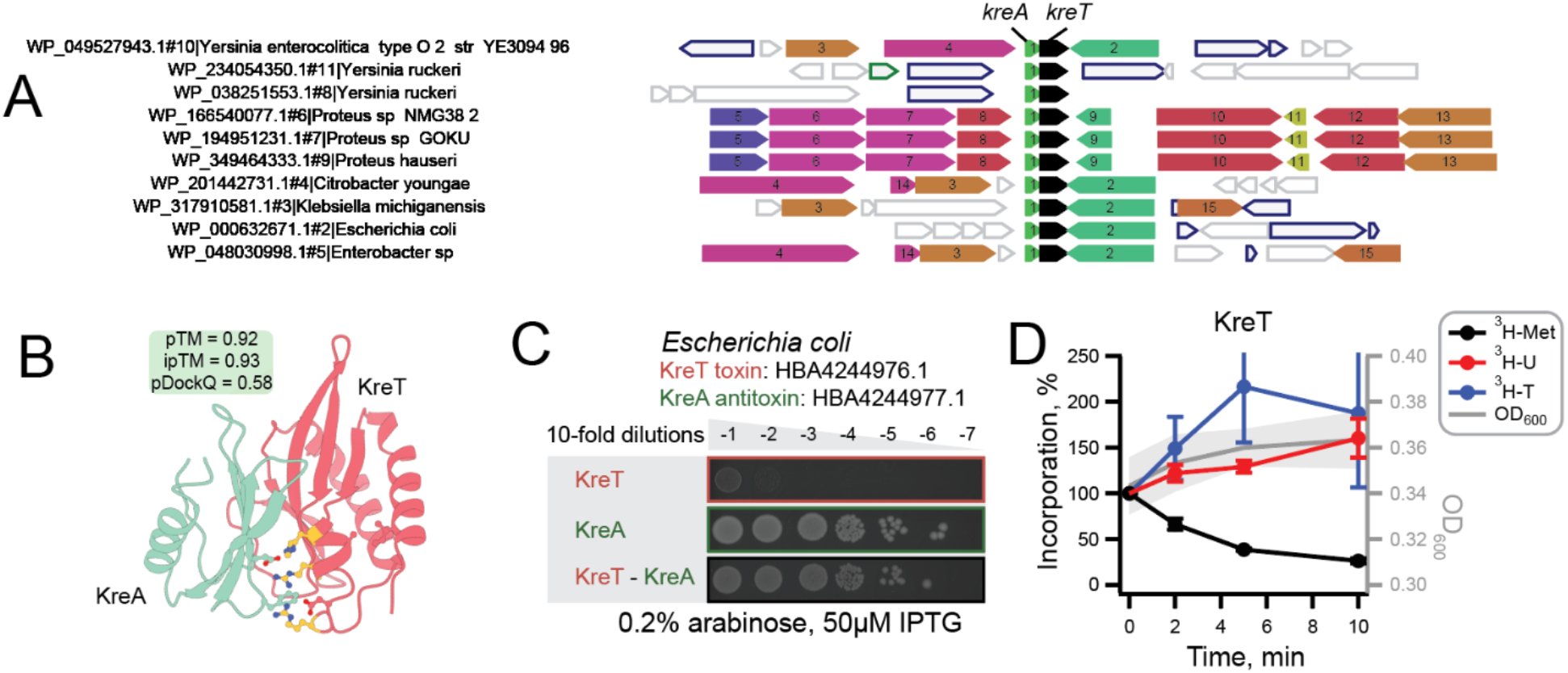
KreTA is a prophage-encoded toxin-antitoxin system. (**A**) Gene neighbourhood analysis identifies putative TA system kreTA encoded in prophage regions of bacterial genomes (visualised here with FlaGs). The kreT gene used as a FlaGs query is shown in black. Proteins encoded by the numbered genes are annotated as follows: 1: KreA; 2: recombinase/integrase; 3: major capsid protein; 4: tape measure protein; 5: deoxyribose-phosphate aldolase; 6: thymidine phosphorylase; 7: phosphopentomutase; 8: purine-nucleoside phosphorylase; 9, 13, 14: hypothetical proteins; 10: pyridoxal-dependent decarboxylase; 11: DksA/TraR family zinc finger protein; 12: MFS transporter; 15: LysR family transcriptional regulator. (**B**) KreA and KreT are predicted to form a dimer. Putative RNase active site residues determined from [35] and interfacing residues are shown as sticks. Interfacing active site residues are colored yellow. (**C**) Expression of the KreT is toxic to E. coli BW25113 cells, but the toxicity is efficiently rescued by the antitoxin KreA. (**D**) Metabolic labelling assays follow the incorporation of ^3^H-methionine (black traces), ^3^H uridine (red), and ^3^H-thymidine (blue) upon expression of KreT.

KreT and KreA proteins are predicted confidently to dimerize (**Fig. 6B**). The proteins are classed as functionally dark in that they belong to communities that have no InterPro domain hits (**Dataset S1**). Foldseek [4] also does not identify any homologous proteins with known function. However, Dali [5] indicates distant but significant fold similarity of KreT to ribonuclease RegB, an endoribonuclease that controls the expression of a multiple phage early genes and which has no identifiable sequence similarity to anything else of known function [35, 36](**Fig. S5A)**. Indeed, the closest structural relative of KreT in VAD is annotated as RegB, but the two proteins are not confidently alignable at the sequence level. In the KreTA dimer, KreA binding sequesters the predicted RNase active site of KreT (**Fig. 6B**), indicative of KreA acting as an antitoxin. Furthermore, Dali suggests KreA may have a broadly similar fold to the immunity protein Tri^Tu^ that neutralises ADP ribosylase toxin Tre^Tu^ [37] (**Fig. S5B**).

To test our prediction that KreTA is a TA, we carried out a toxicity neutralisation assay. Indeed, while expression of KreT is toxic to *E. coli*, this toxicity is efficiently neutralised by the KreA antitoxin, validating that this is TA system (**Fig. 6C**). To investigate the mechanism of KreT toxicity, we performed metabolic labelling using radioactive precursors for translation, transcription and replication. As manifested by selective inhibition of ^3^H-methionine incorporation, KreT inhibits translation, which is characteristic of RNase TA toxins [38] (**Fig. 6D**). Taken together, the fold similarity of KreT to endonucleases and its inhibition of translation suggest it may function by cleaving mRNA, as observed for many known TA toxins [38]. Interestingly, while KreT and KreA are viral in origin, being encoded on prophages, there are no relatives of either in VAD that are identifiable at the sequence level. This highlights another challenge for understanding viral proteome diversity: that viruses can go under the radar by hiding within the genome of their hosts. While ‘cryptic’ infections are most well-known for phages, this is even the case for some eukaryotic viruses [39, 40].

### Conserved enzymatic folds are co-opted in antiviral defence and counter-defence

Viruses are territorial; when they have infected a host cell, they need to keep out competitor viruses. This is called superinfection exclusion and is observed in viruses that infect bacteria as well as eukaryotes [41, 42]. Indeed, temperate bacteriophages have been found to carry a rich diversity of phage defence systems [43]. To analyse defence-like folds in VAD, we used FoldSeek to search our structures against the DefenseFinder database [44].

The largest number of hits to one defence system protein is for the DarG protein from the DarTG system with nine hits to the same hetero-host structural cluster with a eukaryotic host type representative (**Fig. 7A**; pink bar). Proteins in this cluster are all annotated as macrodomain or ADP-ribosyl glycohydrolase proteins (**Dataset S1**). The alignments of these proteins to DarG align solely to their macrodomain, and not to the C-terminal region implicated in binding DarT (**Fig. S6**) [45]. These hits may actually be counter-defence-related via reversal of host ADP-ribosylation catalyzed in eukaryotic antiviral responses [46]. However, since macrodomains have multiple functions, we can not rule out other roles beyond defence or counter defence. Strikingly, six of the structural hetero-host cluster proteins with hits to DarG belong to sequence mono-host clusters, indicating a conservation of macrodomain folds across viruses infecting different domains of life, even when sequence similarity is not apparent (**Fig. 7A**).

**Fig. 7.**
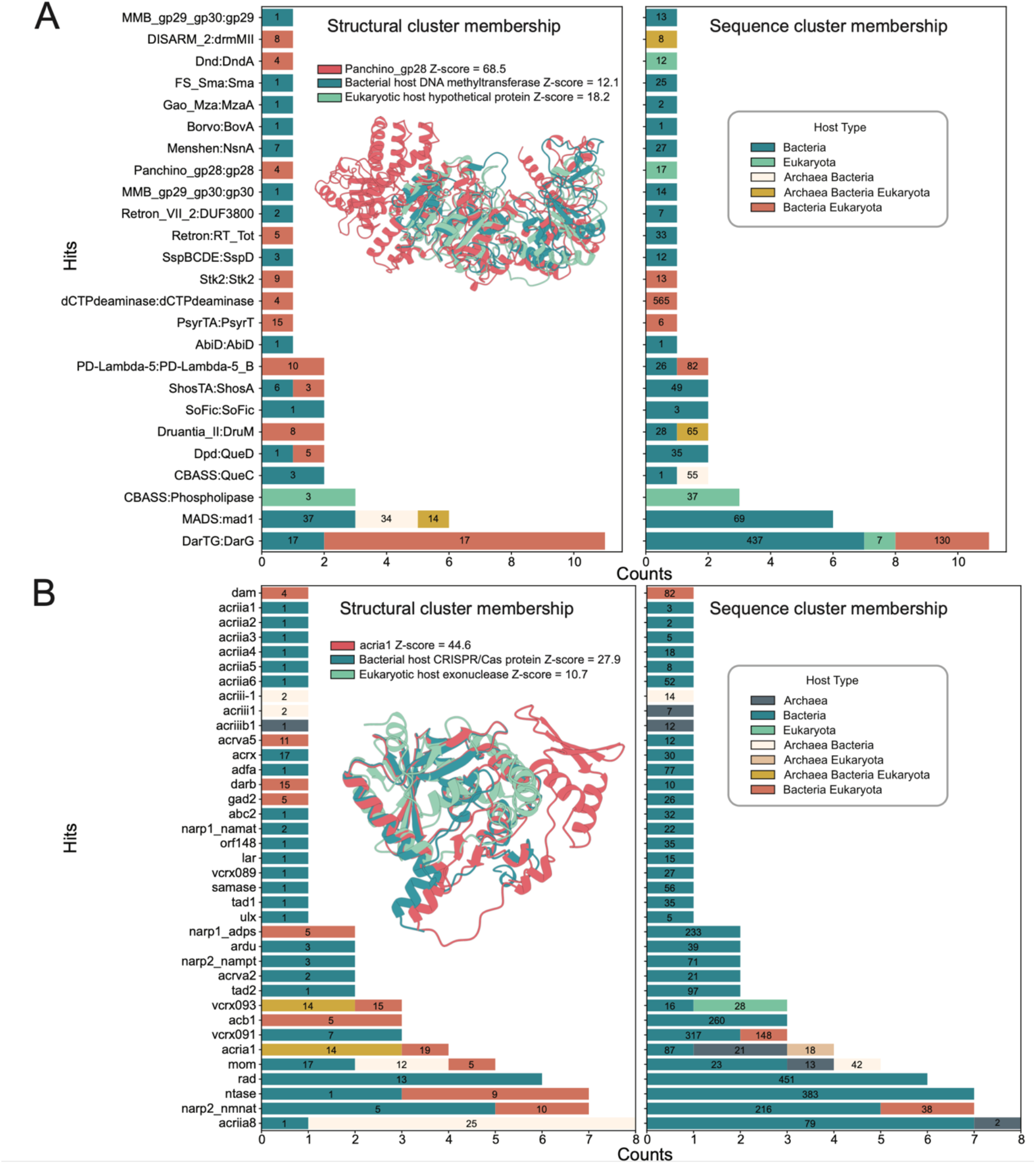
Antiphage defence-related proteins in VAD. Hits are coloured by structural cluster host type, numbered by structural cluster size (left), and by sequence cluster host type and sequence cluster size (right). Each panel includes an example defence system protein, a VAD hit aligned to it, and a protein from a different host type within the same structural cluster, aligned using DALI. (**A**) DefenseFinder structural hits, with the example alignment showing Panchino_gp28 (Z-score = 68.5), a VAD hit (eukaryotic host hypothetical protein, Z-score = 18.2), and a different host type protein (Bacterial host DNA methyltransferase, Z-score = 12.1). Numbers on bars refer to the size of the cluster that the protein with a hit to a defense protein belongs to. For example, DarG has 11 hits to two different cluster host types, each of which has size 17 (**B**) Combined structural hits from AntiDefenseFinder and dbAPIS, with the example alignment showing AcrIA1 (Z-score = 44.6), a VAD hit (phage CRISPR/Cas protein, Z-score = 27.9), and a different host type protein (viral exonuclease (eukaryotic host), Z-score = 10.7).

Another notable hit to defence proteins is that of gp28 in the Panchino system. This protein contains restriction, modification, and specificity domains of type I restriction modification proteins [47]. The VAD protein, which matches Panchino_gp28, is a hypothetical protein belonging to a hetero-host structural cluster with a eukaryotic host-type representative. Within this cluster, the best-aligning protein is a bacterial host-type methyltransferase (**Fig. 7A, Dataset S1**). Proteins in this cluster all have PDB hits to methyltransferase proteins (**Dataset S1**). The alignments to Panchino_gp28 cover mostly the methyltransferase domain but also extend to the restriction domain (**Fig. 7A**). As before with DarG, the VAD protein with a hit to Panchino_gp28 belongs to a sequence mono-host cluster and a structural hetero-host cluster, providing another example of cross domain conservation of fold not apparent on the sequence level (**Fig. 7A**).

As well as defending against superinfection, viruses carry counter defence systems. To investigate whether counter defence folds are found in VAD, we searched against anti-defence proteins in the DefenseFinder database [44] and dbAPIS [48]. The anti-defence protein with the most hits to VAD proteins is AcrIIA8 (**Fig. 7B**), a phage head-tail adapter protein that – surprisingly – is implicated in anti-defence by inhibiting Cas9 activity [49]. AcrIIA8 has seven hits within the same structural cluster containing bacterial and archaeal host-type proteins, along with one hit to a bacterial host-type singleton cluster (**Fig. 7B**). When clustered by sequence, AcrIIA8 hits belong to either archaeal or bacterial host-type clusters, and these clusters merge at the structure level, reflecting a conserved fold across host types (**Fig. 7B**). Proteins in these clusters are similarly annotated as phage head, neck, and tail structural proteins, and it is unclear whether they could also possess additional moonlighting anti-defence functions (**Dataset S1**).

AcrIA1, an anti-CRISPR exonuclease affecting spacer acquisition in CRISPR-Cas-mediated defence [50] has hits to VAD proteins belonging to sequence and structure hetero host clusters (**Fig. 7B**). Structural representatives of these clusters are annotated as CRISPR-Cas associated and/or exonucleases, including a PD-(D/E)XK superfamily nuclease (**Dataset S1**) [51]. AcrIA1 aligns well with its bacterial host-type CRISPR-Cas-associated hit, as well as a eukaryote-host exonuclease found in the same structural cluster (**Fig. 7B**). This eukaryote-host exonuclease is remarkably similar in both sequence and structure to phage exonucleases (**Dataset S1**). Furthermore, it has a hit to another CRISPR-Cas-associated anti-defence protein, VCRX093 [52] (**Fig. 7B, Dataset S1**). Given that eukaryotes do not carry CRISPR-Cas immune systems, this nuclease likely has an as-yet-undefined, role in nucleic acid cleavage during the viral life cycle. In addition, its Foldseek hits to R354—a Cas4-like nuclease from the MIMIVIRE virophage resistance system in mimiviruses [53]—raises the possibility that it may represent a functionally analogous, virophage defence or genome processing element.

Finally, RNA ligase T (LigT)-like phosphodiesterases have been found to be conserved across viruses that infect bacteria and eukaryotes [10].This class of proteins includes the anti-defence protein anti-CBASS protein (Acb1), which degrades cyclic nucleotides [54]. Acb1 also has hits to VAD proteins belonging to a structural hetero host cluster (**Fig. 7B**), further supporting its functional role in cross-domain counter-defence.

The observed conservation in viral defence and counter-defence systems is primarily confined to relatively large enzymes such as methyltransferases, ADP-ribose processing macrodomains, and exonucleases. In contrast, small inhibitors, which are particularly common in counter-defence, appear to be less conserved. However, it is important to note that small proteins are inherently more difficult to detect confidently in structural homology searches, especially if their function depends on – and structure changes upon – oligomerisation. Overall, while viruses employ diverse strategies to evade or counteract host defences, the structural folds of enzymatic proteins involved in these processes are more likely to be preserved – or at least detectably conserved – across different domains of life. Notably, these conserved enzymatic folds are not unique to viruses but are repurposed from universal protein architectures that span across bacteria, archaea, and eukaryota. This structural conservation highlights the reuse of molecular mechanisms in host-virus interactions, underscoring that core and likely ancient protein folds remain essential components of defence and counter-defence systems.

### Conclusions

Comprising high-quality structural predictions of monomers and dimers, the Viral AlphaFold Database (VAD) provides a resource for exploring viral diversity, evolution and biology. There is significant functional “darkness” within viral proteomes, and much novel biology awaiting discovery. Through searching for similarities of viral protein structures to communities of proteins in the dark corners of the protein universe, we have uncovered a novel *Enterobacteriales* prophage-encoded toxin-antitoxin system, KreTA.

Our large-scale structural analysis reveals that viruses, despite their vast sequence diversity, often rely on conserved folds that cross host boundaries and are found throughout the tree of life. The structural similarity of viral proteins and defence-related systems illustrates how enzymatic folds are recurrently co-opted in host-virus conflicts. Collectively, our findings illustrate that viral proteins, despite their immense sequence and structural diversity, often rely on conserved structural frameworks, which play central roles in infection, defence, and adaptation across all domains of life.

## Methods

### Sequence acquisition and host assignment on proteins

647,000 virus protein sequences were downloaded from the NCBI RefSeq [19] database (ncbi.nlm.nih.gov; April 2023). NCBI records for each protein sequence in the RefSeq virus dataset are used to attain taxonomy IDs for the virus the protein is found in. These IDs are then matched to entries in the Virus-Host database [20] to assign a host lineage to each protein. Approximately 10,000 RefSeq virus records lack taxonomy IDs, and about 3,000 virus taxonomy IDs are not found in the Virus-Host database. Proteins associated with these missing or unrecognized taxonomy IDs, as well as those from viruses that infect other viruses and proteins without assigned hosts in the Virus-Host database, are labelled as ‘Host Undefined.’ While these proteins are included in the dataset, they do not contribute to host-type classifications. Through manual curation, certain Virus-Host database hosts were determined to be wrongly or insufficiently classified. Viral lineages *Flyfo siphovirus Tbat1_6, uncultured phage*, and *Leviviridae sp*. were set to bacteria-infecting. Viral lineages *Dunaliella viridis virus* and *Tetraselmis viridis virus* were set to “host undefined”.

### Sequence clustering and host assignment on clusters

MMSeqs2 (v. 15.6f452) [21] was used to cluster RefSeq viral sequences and select representatives for structural analysis. The sequence Identity threshold was set to 30%, and the query to target coverage threshold was set to 10%. This resulted in 117,479 clusters with 61,868 singletons. The clusters were assigned a host type based on the host types of the proteins within them. To limit the effect of possible isolated cases of virus-host missannotation on hetero-host assignment, we set a minimum threshold on the number of mixed hosts in order for a cluster to be assigned a hetero-host label. We required that at least three proteins or 1% of the total cluster size (whichever is bigger) be of different host origin. The remaining clusters are classified as archaea, bacteria, or eukaryota mono-host clusters. Only clusters composed solely of proteins without a defined host are classified as “host undefined”.

### Structural prediction of monomers and homodimers

Protein structure predictions were made with AlphaFold2 v2.3.2 [1] with default parameters. Homodimers were predicted with the AlphaFold-Multimer protocol [55]. The structural template cut-off date was set to May 14, 2020 (--max_template_date=2020-05-14). 63 sequence cluster representatives were not predicted due to out-of-memory and database asynchronicity errors in the AF2 pipeline.

The quality of structural predictions is assessed with predicted local difference test (plDDT) [1] scores for monomers, predicted template-modelling (pTM+ipTM) [55], and predicted DockQ (pDockQ) [56] scores for homodimers. The best models, determined by plDDT scores for monomers and pTM+ipTM scores for homodimers, were selected for further analysis.

Foldseek [4] exhaustive search without thresholds was used to perform all vs. all alignments of VAD monomers.

### Structural clustering and host assignment on clusters

VAD was structurally clustered with Foldseek (v. 8.ef4e960) [4] in TMalign mode. Threshold parameters for the clustering were set at 90% for alignment coverage, 0.4 for TM-score, and 0.001 for E-value. Structural clusters were assigned a host type as was done with sequence clusters.

### Searches against other databases

VAD structures and Foldseek [4] were used to search against the following structural databases: AFDB [12], PDB [11], ProteinUniverseAtlas community representatives [13]. The DALI webserver [57] was also used to search the PDB [11].

### Genomic context analysis

For the 871 viral protein clusters belonging to prokaryotic hosts (i.e., Bacteria, Archaea, and Bacteria and Archaea) with a match to a dark community in the Protein Universe Atlas network (i.e., median functional brightness below 5%) with at least 5 protein members, genomic context analysis was conducted using GCsnap [58] with default parameters. For the 2313 tested Protein Universe Atlas community [13], all UniProt IDs linked to that community were used as input targets. To aid visual inspection of gene neighbourhoods, FlaGs2 [59] was also employed.

Conserved genomic windows identified by GCsnap were analysed for the distribution of protein family names (as automatically assigned by GCsnap) around viral protein targets. To uncover communities containing conserved bicistronic gene arrangements with characteristics of toxin-antitoxin systems, genomic contexts containing target proteins from at least six different species were systematically screened. This approach was based on criteria adapted from NetFlax [35] and further generalized through analysis of GCsnap outputs across diverse datasets. The gene pattern prediction relied on the following criteria: (1) genes within a pattern must appear in the same order; (2) intergenic distances between genes within the pattern should not exceed 100 nucleotides, while distances to conserved genes outside the pattern should exceed 100 nucleotides; (3) neighbourhood conservation should be restricted to the genes within the pattern; and (4) the relative frequency difference between a pattern and the most frequent longer variant (i.e., one additional gene) should be maximal within the genomic context. This screening approach identified the most promising dark communities potentially harbouring toxin-antitoxin systems, providing a focused set of candidates for further inspection and functional analysis.

### Predicting higher order oligostates

Oligomeric states for PDB structures were obtained from the Swiss-Model Template Library [60]. To predict the oligomeric state of a VAD cluster protein, we ran a Foldseek search against all chains in the PDB with a TM-score threshold of 0.5 and returned the most common oligomeric state of all the hits found weighted by TM-score.

### Searching against defence and antidefence databases

DefenseFinder [44] models were downloaded from https://defensefinder.mdmlab.fr/wiki/structure on Jan 16th 2025. For each defence system protein, one monomeric model was used in analyses. Anti-defence protein hidden Markov models were manually downloaded from https://github.com/mdmparis/defense-finder-models commit ca8f119. PyHMMER (v 0.10.15) [61] function *most_probable_sequence* was used to extract sequences from hidden Markov models. Alphafold2 [1] version 2.3.2 was used to model sequences extracted with the same runtime parameters as VAD structure prediction. This dataset is referred to as AntiDefenseFinder. dbAPIS [48] models were downloaded from https://bcb.unl.edu/dbAPIS/downloads/ on Feb 9th 2024. Foldseek search without prefiltering and with TM-score threshold 0.5 was used to compare VAD structures to DefenseFinder, AntiDefenseFinder and dbAPIS models. For each VAD protein the hit with the best identity fraction times query coverage was kept. After exhaustive searching further filters were applied when plotting hits. DefenseFinder: alignment TM score > 0.65, e-value < 0.001, average alignment coverage > 0.6, cluster average TM-score > 0.5. AntiDefenseFinder + dbAPIS, alignment TM score > 0.65, e-value < 0.001 (for AntiDefenseFinder only), average alignment coverage > 0.5, cluster average TM-score > 0.5, intra cluster alignment length > 100. Additionally, when plotting AntiDefenseFinder and dbAPIS hits were combined where the hit with the best TM-score per VAD protein was kept.

### Toxin-antitoxin neutralisation assays

The experiments were performed as described earlier, with minor modifications [13]. The genes encoding the candidate toxin (*kreT* WP_000632671.1) and candidate antitoxin *kreA* (WP_ 001008346.1) were cloned into pBAD33 and pMG25 backbones, respectively, yielding VHP1906 (*kreT*) and VHP1970 (*kreA*) plasmids. *E. coli* BW25113 cells were co-transformed either with the plasmid pair expressing the TA system *in trans*, or with one of the two protein-expressing plasmids being swapped for the appropriate empty vector. The cells were grown for five hours at 37°C with shaking at 200 rpm in LB medium supplemented with carbenicillin (100 μg/mL, to maintain pMG25 and its derivatives) chloramphenicol (25 μg/mL, to maintain pBAD33 and its derivatives) as well as 0.2% glucose for repression of the toxin expression. Next, the cells were diluted in an LB medium to final OD_600_ of 1.0, and serial 10-fold dilutions (from 10^-1^ to 10^-7^) were made in LB. The dilutions were spotted on LB agar plates supplemented with 100 μg/mL carbenicillin, 25 μg/mL chloramphenicol, 50 μM IPTG (for antitoxin expression) and 0.2% arabinose (for toxin expression). The plates were scored after an overnight incubation at 37°C.

### Metabolic labelling assays

The labelling assays were also performed as described earlier, with minor modifications [13]. A single colony *of E. coli* BW25113 cells expressing the KreT toxin from the pBAD33 derivative plasmid under control of arabinose-inducible P_*BAD*_ promoter (VHP1976) was grown overnight in 2 mL Neidhardt MOPS minimal medium [62] supplemented with 1% glucose, 0.1% casein hydrolysate, and 25 μg/mL chloramphenicol at 37°C with shaking at 160 rpm in OLS Aqua Pro Shaking Water Bath (Grant Instruments). The overnight culture was used to inoculate a 20 mL culture in MOPS minimal medium supplemented with 0.5% glycerol as well as a set of 19 amino acids lacking methionine (each at 25 µg/mL) to the final OD_600_ of 0.05.

The cells were grown in a 100 mL flask at 37°C with shaking (160 rpm) in a shaking water bath until recovery after the diauxic shift was observed (OD_600_∼0.2-0.3). At this moment, a 1-mL zero time point aliquots were taken and combined in 1.5 mL sterile Eppendorf tubes with either ^3^H-thymidine (Perkin Elmer) (10 µL per point, 4.5 µCi “hot” and 4.8 μM “cold” nucleotide in autoclaved distilled water), ^3^H-uridine (Perkin Elmer) (10 µL per point, 0.56 µCi “hot” and 4.97 μM “cold” nucleotide in autoclaved distilled water) or ^3^H-methionine (Revvity) (10 µL per point, 0.67 µCi “hot” and 14.99 μM “cold” amino acid in autoclaved distilled water). After an 8-minute incubation at 37°C, the zero time point labelling reactions were quenched by the addition of 200 µL of ice-cold 50% trichloroacetic acid (TCA) and transferred on ice.

Immediately after the collection of the zero time, the KreT expression was induced by adding L-arabinose to the culture to final concentration of 0.2%. At 2, 5, 10 and 15 minutes post-induction, 1 mL culture aliquots were taken, combined with the ‘hot’ label and processed analogously to the zero time samples. OD_600_ values were recorded for each time point. The TCA-quenched samples were filtered through GF/C filters (Whatman) pre-washed with 5 mL 5% TCA. The filters washed with 5 mL (twice for ^3^H-methionine and ^3^H-uridine and three times for ^3^H-thymidine) of ice-cold 5% TCA, and then with 95% ice-cold ethanol (5 mL washes, twice for ^3^H-methionine and ^3^H-uridine, three times for ^3^H-thymidine). The filters were placed in 20 mL scintillation vials (Sarstedt) and air-dried until dry (at least for 2 hours) at room temperature. 5 mL of EcoLite Liquid (MP Biomedicals) scintillation cocktail was added to each vial, followed by shaking for 15 minutes prior to counting. The radioactivity was quantified in CPM (counts per minute) using Hidex 600 SLe automatic liquid scintillation counter (Hidex). CPM values were normalized to OD_600_ at each time point, and incorporation percentages were calculated by dividing the normalized CPM/OD_600_ values by the corresponding value at zero time point values. All experiments were performed using three independent biological replicates and presented as mean values ± SD.

## Data visualisation

Structures were visualized and superimposed with Chimera [63], metabolic caballing data were visualized using Igor Pro 7 (WaveMetrics). The Pavian package [64] was used to plot Sankey diagrams.

## Data availability

VAD and AntiDefenseFinder predicted structures are available at https://data-sharing.atkinson-lab.com/vad/

## Acknowledgements

This work was supported by the Knut and Alice Wallenberg Foundation (project grant 2020-0037 to GCA and VH), the Swedish Research Council (Vetenskapsrådet) grants (2019-01085, 2022-01603 and 2024-06071 to GCA; 2021-01146 and 2024-06059 to VH, the Estonian Research Council (PRG2696 to VH), Göran Gustafsson Foundation for Research in Natural Sciences and Medicine (the Göran Gustafsson Prize to VH), The Royal Physiographic Society of Lund (Endowments for the Natural Sciences, Medicine and Technology, number 45414 to RO), and Erasmus+ traineeship grant to MA.

