## Supplementary material for "The Viral AlphaFold Database of monomers and homodimers reveals conserved protein folds in viruses of bacteria, archaea, and eukaryotes": SI_figures_and_tables

**Supplementary Figures**

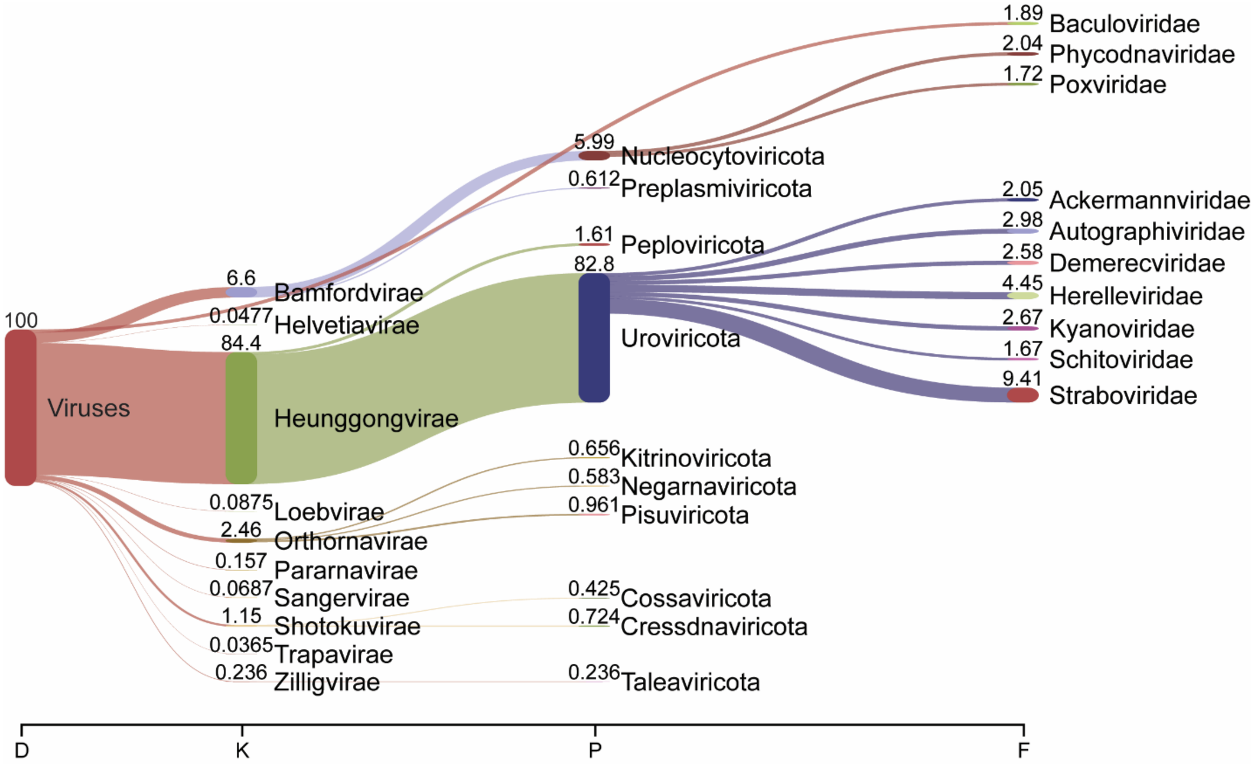


**Fig. S1. Taxonomic distribution of RefSeq viral sequences.**
The ten most abundant taxa per ranks Domain (D), Kingdom (K), phylum (P), family (F), and their relative abundance are shown

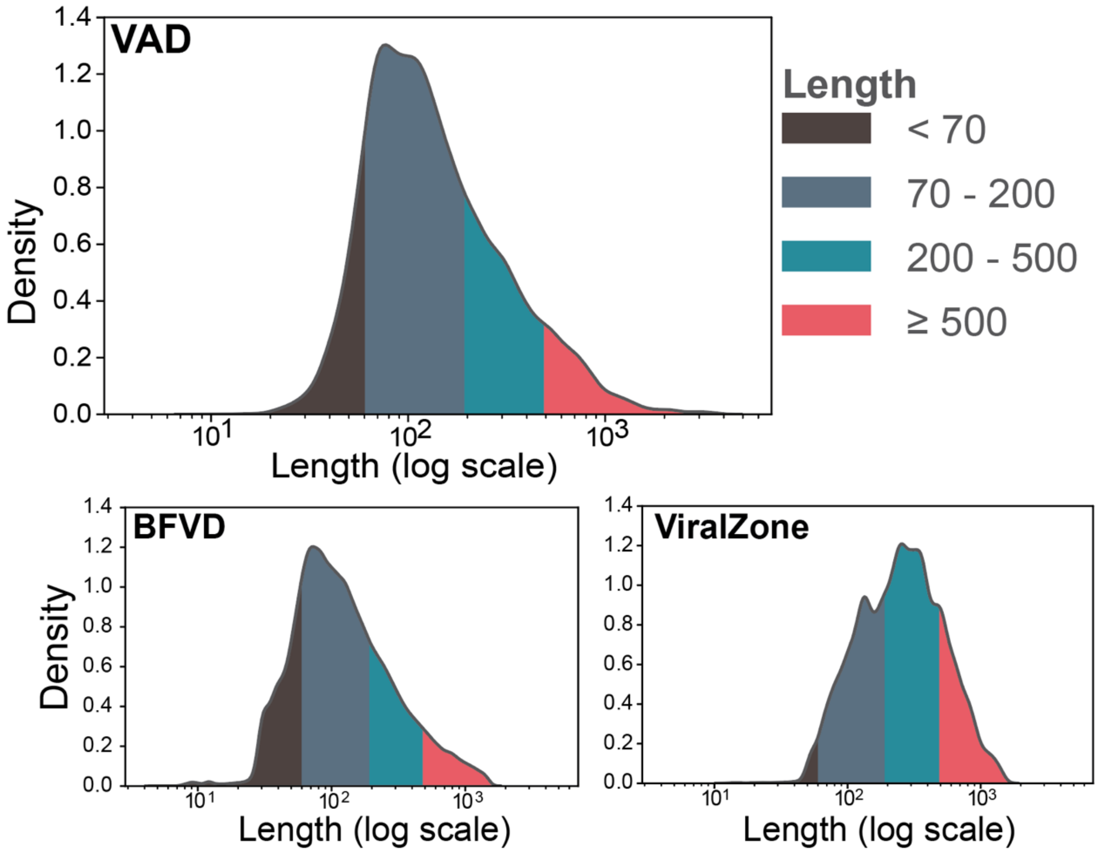


***Fig. S2. VAD comparison with other databases of viral predicted structures**. (**A**) Density plot of Average pLDDT distributions for VAD, BFVD, and ViralZone. pLDDT is averaged per prediction in each dataset. (**B**) UpSet plot detailing overlaps of non-singleton structural clusters (with Foldseek easy-cluster coverage 0.7 and default parameters) across VAD, BFVD, and ViralZone with singleton cluster counts listed (**C**). Three examples of VAD cluster singletons with >80 pLDDT. (**D**). Three examples of clusters unique to VAD with >80 pLDDT. (**E**). Three examples where the VAD structure pLDDT is higher than the corresponding BFVD structure pLDDT.*

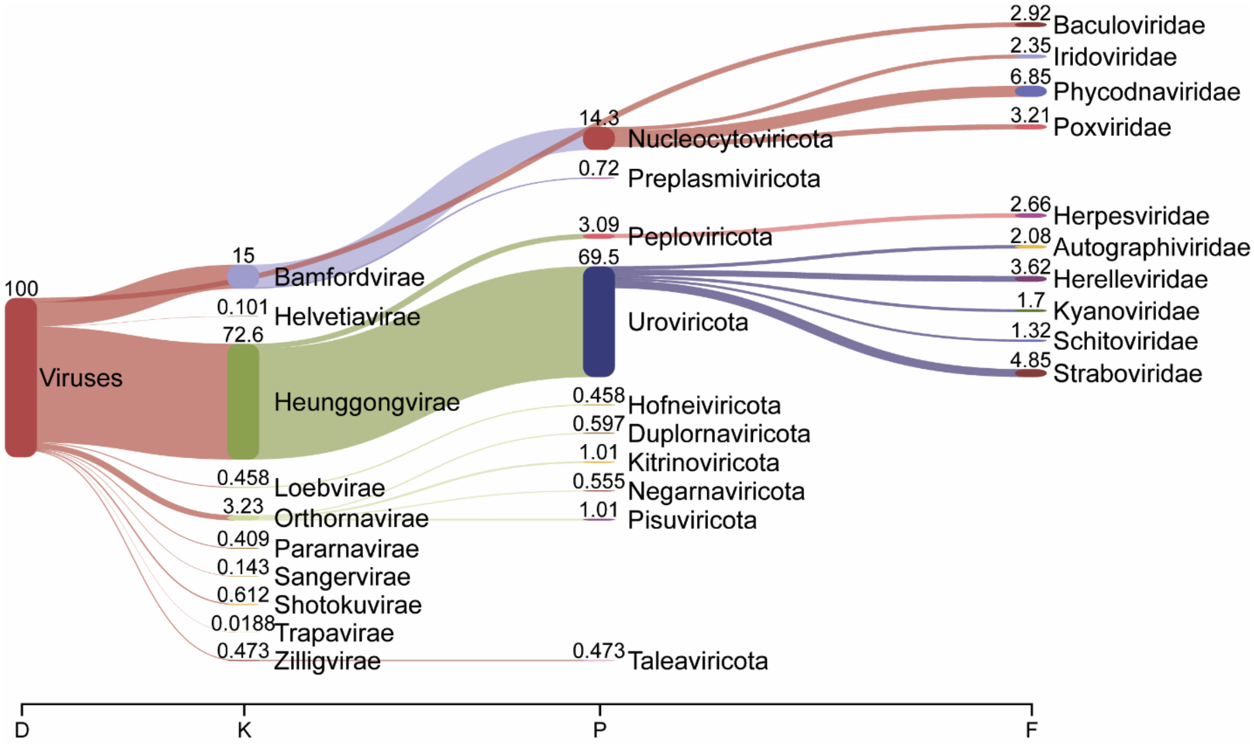


**Fig S3. Taxonomic distribution of VAD.**

The ten most abundant taxa per ranks Domain (D), Kingdom (K), phylum (P), family (F), and their relative abundance are shown.

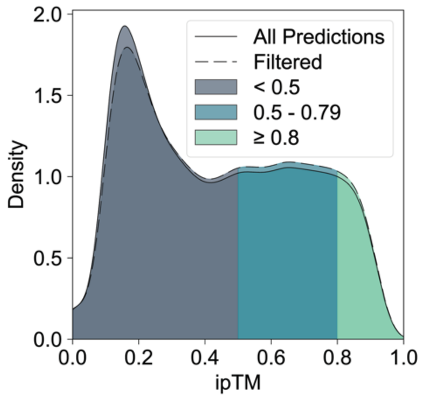


**Fig S4. The ipTM distribution of VAD homodimers does not substantially change when filtered by monomer quality.**

Density plot of ipTM Distribution. Dimeric predictions are filtered based on the average plDDT score of their monomeric counterparts, removing those with an average plDDT below 50.

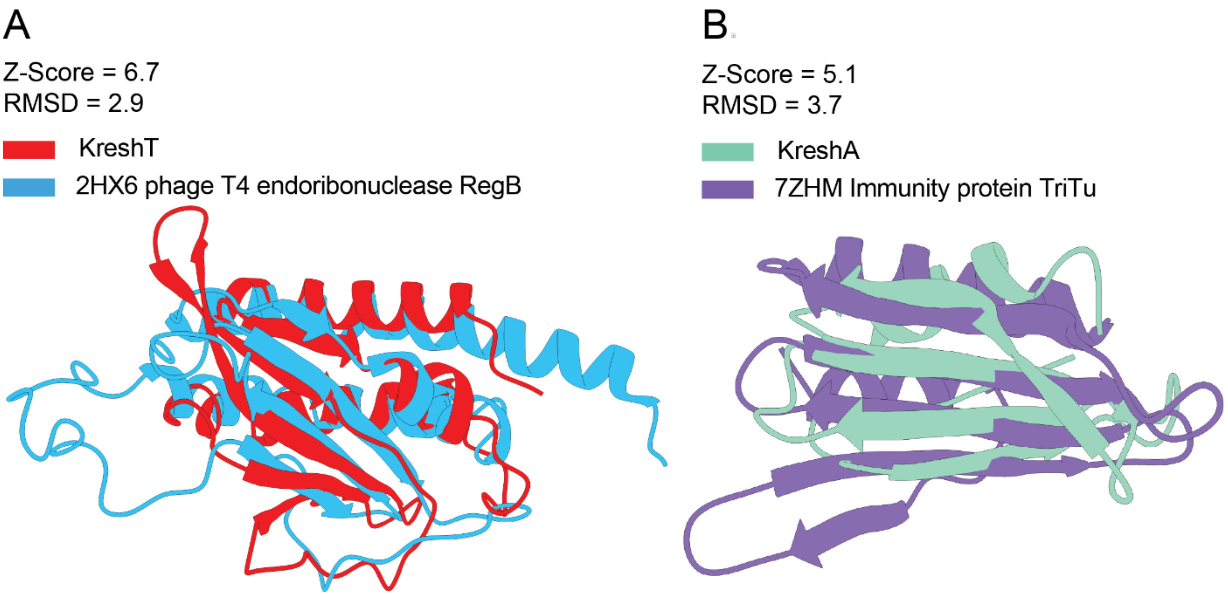


**Fig. S5. Structural search supports KreTA as a novel toxin-antitoxin system.**
(**A**) KreT is likely an endoribonuclease toxin. DALI alignment reveals structural similarity to the phage T4 endoribonuclease RegB (PDB ID: 2HX6), suggesting a potential role in RNA cleavage. (**B**) KreA shows structural similarity to the immunity protein TriTu (PDB ID: 7ZHM), an antitoxin, supporting its annotation as the antitoxin partner in the KreTA system.

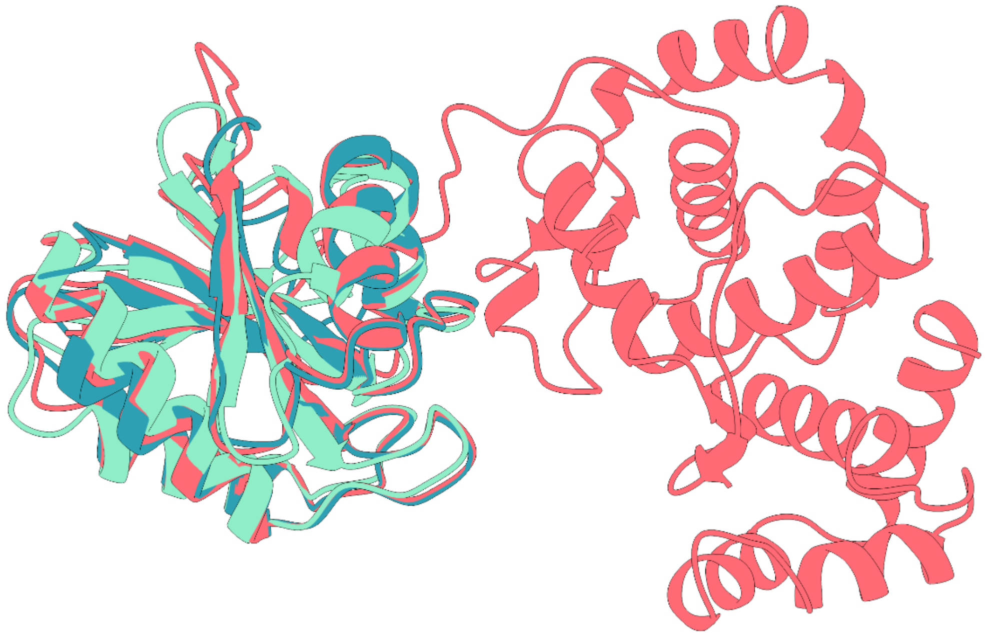

**Fig. S6. DarG partial alignments to VAD proteins, covering macrodomain**.
DarG (red) aligns with eukaryotic host-type (green) and bacterial host-type (blue) VAD proteins within the same structural cluster, showing partial alignment to the N-terminal macrodomain of DarG. Both VAD proteins are hits to DarG.

**Supplementary Tables**

**Table S1. Sequence cluster sizes**

| Sequence clusters | Amount of clusters | Amount of singletons | Amount of Proteins | Average cluster size | Standard deviation of cluster size |
|---|---|---|---|---|---|
| Overall | 117,479 | 61,868 | 647,000 | 5.5 | 18.4 |
| Archaea | 3,465 | 2,243 | 7,061 | 2.0 | 2.3 |
| Bacteria | 86,130 | 42,331 | 521,229 | 6.1 | 19.0 |
| Eukaryota | 25,270 | 15,314 | 99,476 | 4.0 | 14.4 |
| Hetero Host | 388 | 0 | 16,590 | 42.8 | 78.5 |
| Host Undefined | 2,226 | 1,980 | 26,44 | 1.2 | 0.7 |

**Table S2. Structure cluster sizes**

| Structural clusters | Amount of clusters | Amount of singletons | Amount of Proteins | Average cluster size | Standard deviation of cluster size |
|---|---|---|---|---|---|
| Overall | 12,894 | 9,753 | 26,962 | 2.1 | 4.1 |
| Archaea | 163 | 156 | 171 | 1.0 | 0.2 |
| Bacteria | 7,231 | 5,720 | 11,788 | 1.6 | 2.3 |
| Eukaryota | 4,197 | 3,718 | 5,190 | 1.2 | 1.0 |
| Hetero Host | 1,142 | 0 | 9,650 | 8.5 | 10.5 |
| Host Undefined | 161 | 159 | 163 | 1.0 | 0.1 |
